# Range size dynamics can explain why evolutionarily age and diversification rate correlate with contemporary extinction risk in plants

**DOI:** 10.1101/152215

**Authors:** Andrew J. Tanentzap, Javier Igea, Matthew G. Johnston, Matthew J. Larcombe

## Abstract

Extinction threatens many species, yet few factors predict this risk across the plant Tree of Life (ToL). Taxon age is one factor that may associate with extinction if occupancy of geographic and adaptive zones varies with time, but evidence for such an association has been equivocal. Age-dependent occupancy can also influence diversification rates and thus extinction risk where new taxa have small range and population sizes. Here we analysed 509 well-sampled genera from across the plant ToL. We found that a greater proportion of species were threatened by extinction in younger and faster-diversifying genera. Repeating our analyses in two large, well-sampled groups, we found that extinction risk increased with evolutionary age in conifer species but not palms. Potential range size decreased in older, non-threatened conifers more strongly than in threatened taxa, suggesting that range size dynamics may explain differing patterns of extinction risk across the ToL with consequences for biodiversity conservation.

## Introduction

Much of the world’s biodiversity is threatened by extinction because of small geographic ranges and/or population sizes (Pimm et al. 2014). In addition to having traits that promote small ranges and population sizes independent of phylogeny, such as those associated with life history and resource use, some species may be more threatened by extinction because of their evolutionary history (Bennett and Owens 1997; Purvis et al. 2000; Johnson et al. 2002; Arregoitia et al. 2013). Extinction is consequently non-randomly distributed across the Tree of Life (ToL), suggesting that chance events and human activities alone may not be fully responsible for explaining species losses (Bennett and Owens 1997; Purvis et al. 2000; Vamosi and Wilson 2008). Identifying macro-evolutionary predictors of extinction risk can therefore help to assess future conservation status where range and population data are lacking and identify reasons for its non-randomness across the ToL (Jetz and Freckleton 2015).

Taxon age is one measure of the amount of environmental and evolutionary change that species have experienced and may be associated with extinction risk for at least two reasons. The first relates to the idea that older taxa should be less at risk of extinction because they have had more time to disperse across a greater range (Paul et al. 2009; Ceolin and Giehl 2017), consistent with the age-and-area hypothesis (Willis 1926). While differences in the time for dispersal may weaken over long time scales (i.e. millions of years), younger taxa may also face less available space and resources as niches fill through time irrespective of dispersal ability (Tanentzap et al. 2015). As new taxa initially tend to have small range and population sizes, especially if speciation started from small reproductively isolated populations that occupy narrow adaptive spaces (Valente et al. 2010; Castiglione et al. 2017), younger species in rapidly diversifying clades should face a greater risk of extinction (Davies et al. 2011; Greenberg & Mooers 2017).

A second historical explanation for variation in extinction risk relates to differences in niche breadth among species of different ages. Older species may have survived long-term environmental changes because they are more generalist (Liow 2007). As broader niches are positively associated with larger ranges (Slatyer et al. 2013), this explanation would result in another positive age-and-area association. By contrast, there may be a negative correlation between age and extinction risk if older species are more specialised and have smaller ranges. We term this idea the evolutionary specialism hypothesis. Older species can appear more specialised because traits that were once advantageous became less adaptive as environments diverged from past selection regimes (Wilson 1959; Žliobaitė et al. 2017). More specialist species with narrower niches and geographic ranges may only persist over long time periods in refugia or by having large local population sizes (Williams et al. 2009).

The potential for species to expand their range and reduce extinction risk with time may ultimately depend on their mode of speciation. Repeated range expansion and contraction (i.e. “taxon cycles”) that isolate peripheral populations consistent with centrifugal or peripatric speciation can produce small ranges in descendent taxa (Gaston 1998). Consequently, older species may have a lower extinction risk because they have had more time to disperse and expand their range, and experience less niche pre-emption from earlier evolving competitors (Tanentzap et al. 2015). Lineages with high diversification rates under this mode of speciation can similarly face greater extinction by producing species that have small ranges (Schwartz and Simberloff 2001). By contrast, any signature of time in extinction risk distributions may be absent with vicariant speciation because asymmetry in the ranges of ancestors and daughter species is consistently smaller and ancestral species often disappear via cladogenesis (Gaston 1998).

Evidence that taxon age is associated with extinction varies among lineages, so testing across different divisions in the ToL. Previous work in birds (Gaston and Blackburn 1997) and marsupials (Johnson et al. 2002) found that older lineages were more threatened by extinction, whilst the reverse was shown across non-lemur primates (Arregoitia et al. 2013). The only study on plants, to our knowledge, found a higher extinction risk in younger, rapidly diversifying clades of the South African Cape (Davies et al. 2011). Broader generalisations across plants have not been possible until now because of poor taxonomic sampling that prevents reliable divergence times from being estimated.

Here, we tested whether younger and faster-evolving lineages were associated with greater extinction risk across 509 genera representing 9,174 species. We did so by combining the largest time-calibrated phylogenetic tree presently estimated for vascular plants with all available peer-reviewed assessments of conservation status from the International Union for Conservation of Nature (IUCN) *Red List* (2016). We complemented our findings with analyses for two large, ancient, and widespread plant clades (conifers and palms). These analyses allowed us to address concerns around estimating divergence times from the larger but under-sampled phylogenetic tree and threat status from incompletely sampled genera. By working at the species-level, we could also collate geographic distribution data to test the age-and-area and specialism hypotheses, and how they might explain differences in age-extinction correlations between taxonomic groups with contrasting histories. Positive correlations between taxon age and range size would support the idea that older species have had more time for dispersal (i.e. age-and-area hypothesis), whereas a negative correlation would support the idea that older species are more specialist.

## Methods

### Data assembly

We first selected genera for which we could confidently estimate the time of divergence from their sister genera, i.e. ‘stem age’. Genera were selected from the time-calibrated, species-level phylogenetic tree of Qian and Jin (2016), which was an updated version of Zanne et al. (2014). The selected genera came from densely sampled clades (i.e. families) to circumvent low sampling across the broader tree both at a species- and genus-level. For each family, we calculated the proportion of genera that were sampled in the phylogeny from the taxonomic database curated by *taxonlookup* v1.1.1 (Pennell et al. 2016) in R v3.2 and retained those with ≥60% coverage. We also used stem ages because they only require one species to be sampled within each genus and reflect the entire evolutionary history of clades unlike crown ages that can have young age biases because they consider only extant species (Scholl and Wiens 2016). Taxa outside of an established “core clade” for each genus, as determined using *MonoPhy* in R (Schwery and O’Meara 2016), were removed prior to all calculations.

After calculating ages from the large tree, we intersected the selected genera with 25,452 IUCN assessments and calculated the proportion of species in each genus threatened with extinction. Threat status is jointly determined from abundance, recent temporal change in population size, and various measures of geographic distribution, such as occupancy and fragmentation (IUCN 2016). Therefore, metrics of range size alone may not entirely predict extinction risk despite the potential to use these terms interchangeably. We further restricted our analysis to genera with >1 species, of which ≥20% had sufficient data to be assessed for extinction risk. We excluded 154 monotypic genera because these would confound our analyses as they all had the same diversification rate irrespective of lineage age. Overall, 509 genera had both reliable age and risk status data spanning 4,925 IUCN species-level assessments.

We also estimated net diversification rates for the 509 genera. We used a well-over time within genera given a known stem age and species richness (Magallon and Sanderson 2001). Following standard practice, we assumed three values of relative extinction *ε* of 0.0, 0.5 and 0.9 when estimating diversification (Magallon and Sanderson 2001). All taxonomy was standardised to The Plant List nomenclature using the *Taxonstand* R package prior (Cayuela et al. 2012).

We also repeated our diversification analysis as above with two large clades that were well sampled at a species-level in separate time-calibrated phylogenies. These clades included 70% of all 651 accepted Pinales (extant conifers) (Leslie et al. 2012) and all 2,539 Arecaceae (palms) (Faurby et al. 2016). We intersected risk statuses of the two clades with species stem ages, giving *n* = 433 and 547, respectively. For the palms, we used the maximum clade credibility tree that we computed from the posterior distribution of trees that was generated using topological constraints based on Govaerts taxonomy recommended in Faurby et al. (2016).

Finally, we assembled range data for our two large clades. Georeferenced records with no flagged issues were downloaded from the Global Biodiversity Information Facility (www.gbif.org). Conifer data were supplemented by published records absent from GBIF (table A1). All duplicate and spatially invalid records (e.g. non-numerical, exceeding global extent, located in the ocean, urban areas, or country centroids) were removed with the R package *sampbias*. Using the occurrences, we estimated potential range size with a mechanistic species distribution model (SDM) that predicted the physiological tolerances of species for growth from distribution data (Higgins et al. 2012). Absence points for the SDM were generated using standard approaches (details given in Appendix A). We then summed the total number of equal-area (Mollweide projected) 0.25 decimal degree grid cells occupied by each species. We found no evidence that sampling varied systematically with species age in a way that would bias our subsequent analyses (table B1).

### Statistical analyses

We separately tested whether genera with a greater proportion of threatened taxa were correlated with younger ages and faster diversification rates using phylogenetic least squares (PGLS) regression. Although the least squares model assumed normally distributed errors, and the response variable was a proportion with binomial errors, PGLS is appropriate for testing the null hypothesis of no statistically significant effect of an independent variable on a non-Gaussian response (Ives 2015). We also fitted the PGLS regression using the *gls* function in R because this approach, unlike other model fitting functions that incorporated phylogenetic information (e.g. *phyloglm*), could account for different sample sizes across genera by weighting observations with the inverse square-root of the number of IUCN assessments that they received (Garamszegi and Møller 2010). Following standard practice, the PGLS was fitted with maximum-likelihood transformations of branch lengths based on the strength of phylogenetic covariance estimated by Pagel’s *λ* (Orme 2013). Both ages and diversification rates were log-transformed. Models were not fitted with both predictors simultaneously as they were highly correlated (Spearman’s *r* < -0.79). We repeated this analysis in conifers and palms, and again did not simultaneously fit age and diversification rates given high correlations (*r* = -0.78 to -0.91). Fit of the PGLS was summarised by the correlation coefficient *r* between predicted and observed values.

For conifers and palms, we also tested whether extinction risk was associated with younger species and how this was influenced by range dynamics. We first fitted logistic regression models to threat status as a function of species age using penalised maximum-likelihood and accounted for phylogenetic non-independence of species with the *phylolm* R package (Ho et al. 2014). Predictors were scaled to a mean of 0 and standard deviation of 1 to compare effects.

We also tested how potential range size was associated with species age in both conifers and palms. First, we used PGLS to test whether older ages correlated with larger range sizes, which by definition reduce extinction risk (IUCN 2016), and allowed the effect to vary with threat status (i.e. statistical interaction). We expected threatened species would, by definition, always have relatively small ranges, producing an invariant or weak age-range association. By contrast, non-threatened species should reach larger ranges with time if the age-and-area hypotheses was supported, whereas the reverse could be expected under the specialism hypothesis. One limitation with this analysis is that it does not compare threatened and non-threatened species of the same age, and so can introduce biases if there are systematic differences in the ages of these two groups.

To further analyse how potential range size was associated with species age, we undertook a second comparison that focused on pairs of sister species with contrasting threat status. For each pair, we calculated the difference in potential range size between the sisters, so as to avoid pseudoreplication, and correlated this with their age. We compared this association to when sisters had the same threat status to test the null hypothesis that being threatened with extinction does not change age-range associations. Focusing on sister pairs was desirable because it can minimize factors that confound age-range associations, such as unobserved extinctions (Hodge and Bellwood 2015). Range differences can also shed light on the underlying mode of speciation. For example, there may be greater disparity in the ranges of young species pairs under peripatric as opposed to allopatric speciation (Gaston 1998; Hodge and Bellwood 2015), resulting in a negative correlation between age and range asymmetry (fig. A1). This pattern may ultimately result in either a positive or negative association between age and extinction status, depending on whether species expand their ranges with time (i.e. age-and-area hypothesis) or contract their ranges as environments change (i.e. specialism hypothesis). We tested if this correlation between age and range asymmetry was different from randomly sampling the same number of sister pairs 1,000 times, but choosing those where both members of the pair had the same threat status. We chose both species to be non-threatened for conifers and both to be threatened for palms as most identical species pairs in the two clades fell into these two categories (70/85 and 51/68, respectively). Reassuringly, potential range size of non-threatened conifers and threatened palms did not differ in our analysis when sister species had the same threat status, supporting their use as “control” contrasts (t-test: *t*_177_ = 0.183, *p* = 0.855 and *t*_124_ = 0.597, *p* = 0.552, respectively). R code to perform our analyses is in Data S1.

## Results

We found that relatively more species were threatened with extinction in faster diversifying genera (for *ε* of 0.0, 0.5, 0.9: *t*_507_ = 3.64, 3.73, 3.83, respectively; *p* < 0.001 and *r* = 0.15 for all). The mean proportion of a genus threatened with extinction more than doubled from 36% to 84% between the slowest and fastest diversifying genera (fig. 1a). Although these results could have arisen because faster diversifying genera were younger (fig. 1b), as genus age was negatively associated with risk status (*t*_507_ = -2.82, *p* = 0.005, *r* = 0.14), diversification rate was not a simple proxy for age as it had larger effect sizes. A caveat is that we did find some bias in our dataset. Sampled genera were older and slower diversifying, on average, than obtained by applying our sampling criteria to the initial tree (i.e. before intersecting with threat status; table B2). Repeating our analyses with only the genera from the more complete conifer and palm species-level datasets was also inconclusive (table B3), potentially because of small sample sizes (*n* <70; fig. B1). Many conifer genera were also highly threatened despite being old and slowly diversifying (fig. B2).

**Figure 1.**
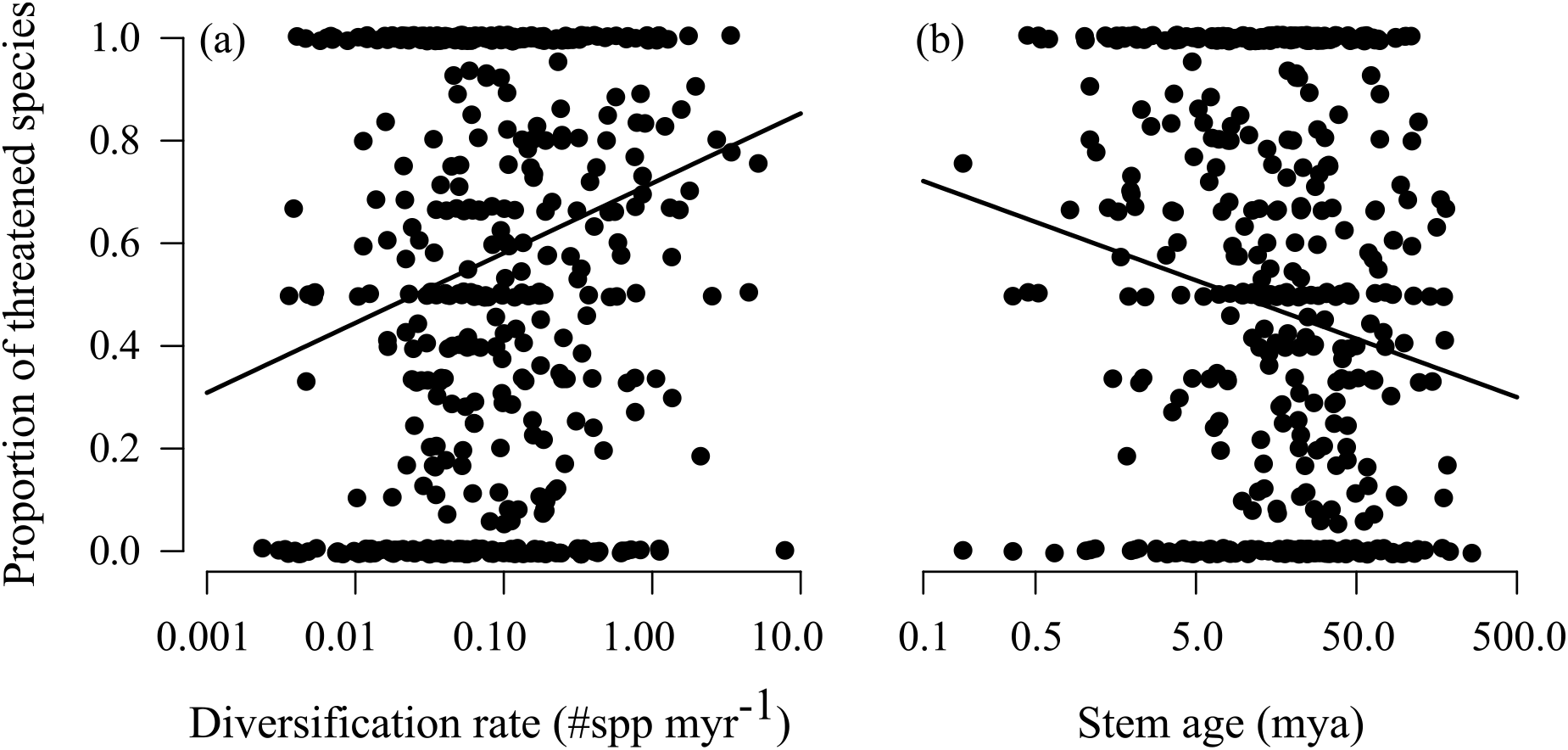
More species are threatened with extinction in (**a**) faster diversifying and (**b**) younger genera. Diversification was estimated for *ε* = 0.50. Solid lines are mean associations estimated by PGLS.

In contrast to our finding across the plant ToL, analyses with the more complete species-level datasets revealed that older conifers but not palms were relatively more threatened by extinction (*z*_431_ = 2.17, *p* = 0.030 and *z*_545_ = -1.70, *p* = 0.089, respectively; fig. 2a). The absolute mean effect ± SE was nearly double in the conifers (0.27 ± 0.12 vs -0.14 ± 0.08 on log-scale), leading to a 31% absolute increase in the probability of being threatened over the range of observed ages (fig. 2b).

**Figure 2.**
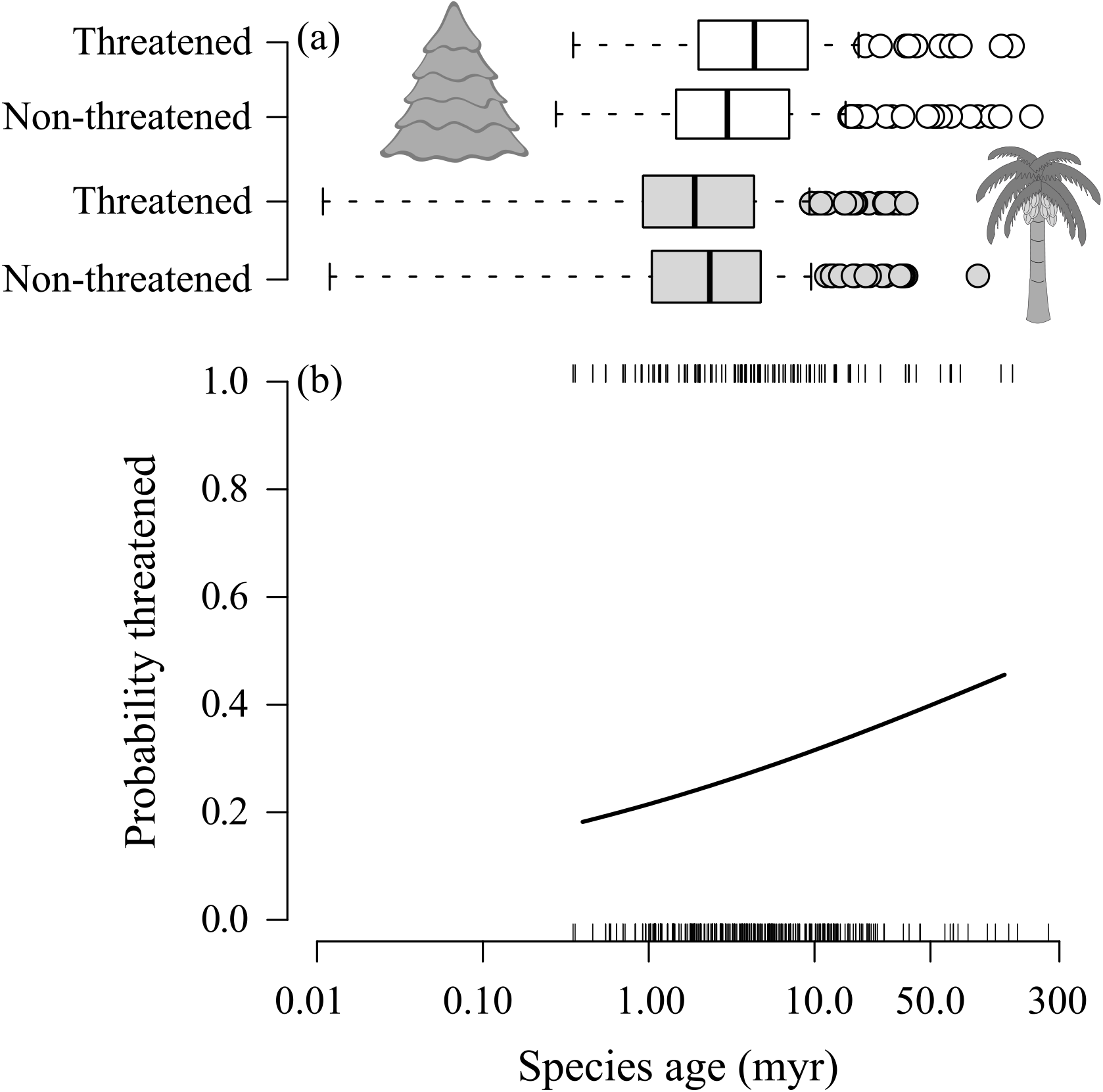
Older conifers but not palms have a greater probability of being threatened by extinction. (**a**) Boxplot for stem ages of conifer (white, *n* = 433) and palm (grey, *n* = 547) species that were classified as either threatened or non-threatened. Solid line is median, box is inter-quartile range, whiskers extend 1.5-times the interquartile range, and points are outliers. (**b**) Change in probability of a conifer being classified as threatened with species age. Solid line is mean association estimated by phylogenetic logistic regression.

A smaller potential range size increased the extinction risk of older conifers, supporting the specialism hypothesis. We specifically found that non-threatened conifers had narrower ranges as their age increased relative to sister species that were threatened (fig. 3); ranges in neither threat status independently changed with age (table B1). As the age of conifers increased, this difference between sister-species pairs of contrasting threat status was larger than expected if sisters had the same threat status (*r* = -0.27, *p* = 0.025; fig. 3a). Contrasting threat status did not alter correlations between age and potential range size in palms, consistent with the lack of an age-extinction association (*r* = -0.14, *p* = 0.222), and there was no correlation between species age and absolute range size (table B1). Larger potential ranges did, however, always reduce extinction risk (table B1). Our results with conifers and palms were also not simply an artefact of biased sampling as ages and rates did not markedly differ from observations across entire clades, i.e. before filtering with IUCN data (table B4).

**Figure 3.**
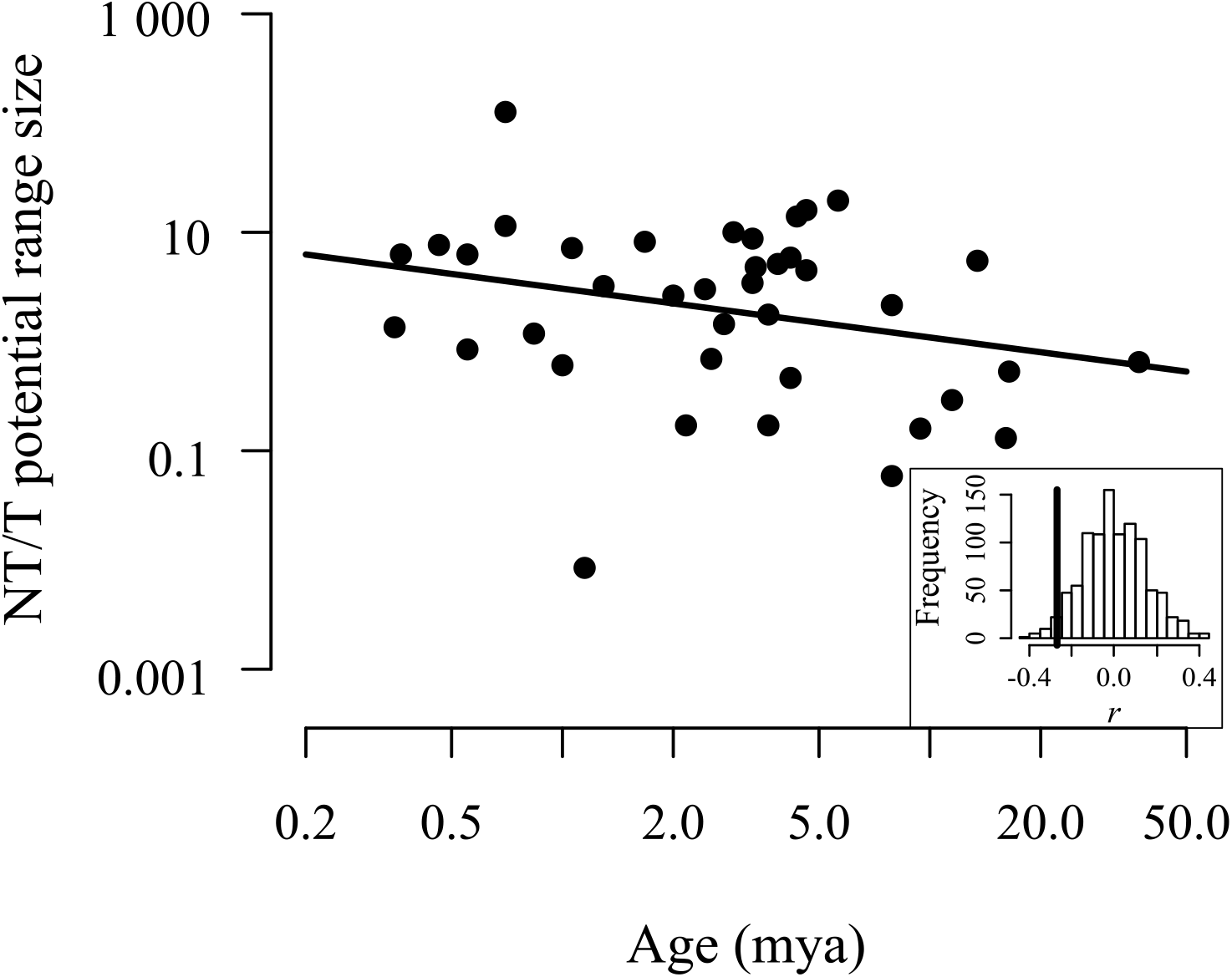
Differences in range size between sister conifers of contrasting threat status decrease with their age. For each sister pair of non-threatened (NT) and threatened (T) taxa for the corresponding correlation coefficient *r*. Inset shows frequency distribution of *r* calculated for 1,000 random simulations of sister pairs of the same threat category, with vertical line denoting observed correlation for contrasting threat status, i.e. corresponding to plotted data points.

## Discussion

Our results implicated range size as a proximate explanation for why clade age and diversification rate were associated with extinction risk in plants. Although our findings across the wider plant ToL contrasted those in conifers, they were consistent with the age-and-area hypothesis in at least two ways. First, young species tend to occupy narrower geographic and adaptive spaces (Castiglione et al. 2017), particularly as most plant speciation involves vicariance (Davies et al. 2011; Anacker and Strauss 2014; Igea et al. 2015). Time may consequently be required for post-speciation range expansions despite much of the available area remaining favourable for establishment (Pigot et al. 2010; Pigot and Tobias 2013; Anacker and Strauss 2014). Second, if species diversification is density-dependent, such as because of limited resources, then younger lineages that occupy smaller ranges will tend to leave more niche space available for young species (Rabosky and Hurlbert 2015). The consequent increase in rates of species diversification will again elevate extinction risk in younger lineages if reproductive isolation arises within small geographic and adaptive spaces. Time-dependent range expansions may be unnecessary under other modes of speciation, e.g. parapatry or sympatry (Pigot et al. 2010), and if range expansion is not limited post-speciation (Schurr et al. 2007). These differences in modes of speciation can also help explain the lack of consistent evidence for age-dependent extinction across the large taxonomic scale in our study and across animals (Gaston and Blackburn 1997; Johnson et al. 2002; Davies et al. 2011; Arregoitia et al. 2013; Greenberg and Mooers 2017).

The global status of conifers differs from palms and other plant clades, potentially explaining why older species had smaller potential ranges that made them more threatened by extinction. Conifer species are older on average than the rest of the Qian and Jin (2016) tree (Welch’s *t*-test: *t*_465.5_ = 13.71, *p* < 0.001), and many species are range-restricted (Farjon 1996; Jordan et al. 2016). Consistent with the evolutionary specialism hypothesis, most old conifers evolved during warmer wetter climates, where they occupied larger ranges than in the present day (Farjon 1996; Jordan et al. 2016). Old species may have only escaped extinction by inhabiting climatic refugia that have been historically stable (Leslie et al. 2012; Condamine et al. 2017). Cycadales, which are closely related to conifers, have undergone similar range contractions because of global cooling, resulting in presently high extinction risk (Yessoufou et al. 2017). By contrast, most palm species have occupied relatively large areas of stable habitat since the Eocene (Kissling et al. 2012), potentially explaining the lack of age-range correlations. Speciation in palms may have also occurred largely by long-distance dispersal (Baker and Couvreur 2013), which can produce less range asymmetry (Gaston 1998). Consequently, palms may lack age-range associations that influence extinction risk. We also cannot exclude the possibility that palm species that were susceptible to environmental change have already gone extinct or traits that make them more prone to extinction are not taxonomically conserved, resulting in no signature of taxon age on extinction (Arregoitia et al. 2013).

Our findings suggest that macro-evolutionary dynamics have some value for biodiversity conservation. Specifically, we found that these dynamics provided an indicator of contemporary extinction risk that might be easier to derive for large numbers of taxa than detailed species-level assessments. Macro-evolutionary dynamics might also offer insight into the vulnerability of species to future change, as the smaller population and range sizes that make some species prone to extinction are likely to be carried into the future (Condamine et al. 2013). Although our results must be interpreted with caution, given biases inherent to our datasets, they provide new evidence that lineages span a continuum from little species turnover to producing fast diversifying and extinction-prone taxa (Greenberg and Mooers 2017). The effect of age that we found at different taxonomic scales also suggests similar patterns should emerge when the plant Tree of Life becomes more densely sampled.

## Data Accessibility

All relevant data and R code to carry out the analyses described in this paper are available at https://github.com/atanzap/age-extinction-plants.

## Acknowledgements

We thank G. Tanentzapf for the question that inspired this study and G. Jordan for sharing the conifer phylogeny. A Mooers, D. Greenberg, and one anonymous reviewer provided comments that improved an earlier version of this manuscript. This preprint has been reviewed and recommended by Peer Community In Evolutionary Biology (https://dx.doi.org/10.24072/pci.evolbiol.100058).

